# Neural selectivity for visual motion in macaque area V3A

**DOI:** 10.1101/2020.09.15.298349

**Authors:** Nardin Nakhla, Yavar Korkian, Matthew R. Krause, Christopher C. Pack

## Abstract

The processing of visual motion is carried out by dedicated pathways in the primate brain. These pathways originate with populations of direction-selective neurons in the primary visual cortex, which project to dorsal structures like the middle temporal (MT) and medial superior temporal (MST) areas. Anatomical and imaging studies have suggested that area V3A might also be specialized for motion processing, but there have been very few studies of single-neuron direction selectivity in this area. We have therefore performed electrophysiological recordings from V3A neurons in two macaque monkeys (one male and one female) and measured responses to a large battery of motion stimuli that includes translation motion, as well as more complex optic flow patterns. For comparison, we simultaneously recorded the responses of MT neurons to the same stimuli. Surprisingly, we find that overall levels of direction selectivity are similar in V3A and MT and moreover that the population of V3A neurons exhibits somewhat greater selectivity for optic flow patterns. These results suggest that V3A should be considered as part of the motion processing machinery of the visual cortex, in both human and non-human primates.

**Significance statement:** Although area V3A is frequently the target of anatomy and imaging studies, little is known about its functional role in processing visual stimuli. Its contribution to motion processing has been particularly unclear, with different studies yielding different conclusions. We report a detailed study of direction selectivity in V3A. Our results show that single V3A neurons are, on average, as capable of representing motion direction as are neurons in well-known structures like MT. Moreover, we identify a possible specialization for V3A neurons in representing complex optic flow, which has previously been thought to emerge in higher-order brain regions. Thus it appears that V3A is well-suited to a functional role in motion processing.

## Introduction

The primate visual system consists of numerous brain regions that represent stimuli in contralateral visual space. Classically, these brain regions have been identified with parallel pathways: a dorsal pathway for spatial vision and a ventral pathway for object vision (Mishkin et al., 1983). Along each pathway, the brain processes information in hierarchical stages, with each stage generating selectivity for more complex stimuli.

Among these brain regions, area V3A has been somewhat controversial, both in terms of its connection to the two pathways and its station along the functional hierarchy. Anatomical studies have often yielded inconclusive results, with some showing that it projects primarily to the dorsal pathway (Ungerleider and Desimone, 1986; Baizer et al., 1991; Felleman and Van Essen, 1991), others indicating that it projects primarily to the ventral pathway (Felleman and Van Essen, 1983; Distler et al., 1993), and a few placing it in both pathways (Morel and Bullier, 1990). From a functional perspective, human imaging studies have frequently reported that area V3A is concerned with motion processing (Tootell et al., 1997; Smith et al., 1998; Sunaert et al., 1999; Braddick et al., 2000; Seiffert et al., 2003; Koyama et al., 2005; Cardin et al., 2012; Furlan and Smith, 2016), while similar neuroimaging studies in monkeys have revealed relatively weak responses to motion in this area (Tolias et al., 2001; Vanduffel et al., 2001; Vanduffel et al., 2002). The few single-unit recordings of motion selectivity in V3A have similarly revealed a paucity of direction-selective neurons (Essen and Zeki, 1978; Gaska et al., 1988; Galletti and Battaglini, 1989), with most neurons being reportedly tuned for static orientation (Essen and Zeki, 1978; Galletti et al., 1990).

One explanation for this discrepancy is that area V3A plays different roles in humans and monkeys (Tootell et al., 1997; Orban et al., 2003; Orban et al., 2004; Sereno and Tootell, 2005), but the existing data on single-neuron direction selectivity in V3A are scant. Gaska et al. (1988) reported that approximately 30% of V3A cells were direction selective, when tested with moving bars, and a similar figure was reported based on qualitative estimates by Galletti et al. (1990). However, both studies reported data from relatively small samples of neurons, tested with a very restricted set of motion stimuli. Given the evidence for functional variation within extrastriate areas like V2 and V4 (Livingstone and Hubel, 1987; Li et al., 2013), and the possibility that V3A encodes a variety of different motion stimuli (Koyama et al., 2005; Furlan and Smith, 2016; Strong et al., 2017), the possibility remains that V3A plays a role in monkeys similar to that proposed in humans.

In this study, we have recorded from a large population of single neurons in macaque V3A. For comparison, we simultaneously recorded from neurons in the middle temporal (MT) area, which is well-established as a motion processing region. When tested with drifting gratings or random dot fields, V3A neurons in our sample exhibited direction selectivity that was, on average, very similar to the direction selectivity in MT. Moreover, when tested with more complex motion patterns (optic flow), many V3A neurons showed strong direction selectivity, and this type of selectivity was more prominent than what was found in MT. Together with the evidence from human imaging (Koyama et al., 2005; Furlan and Smith, 2016; Strong et al., 2017), our results suggest that area V3A contains at least some domains that are highly selective for visual motion.

## Materials and Methods

### Electrophysiological Recordings

Two adult rhesus monkeys (*Macaca mulatta*; 7 kg female and 18 kg male) were used for electrophysiological recordings. An MRI-compatible titanium head post was implanted on each monkey’s skull in order to stabilize their heads during experimental sessions. Eye movements were monitored during experiments with an EyeLink 1000 infrared eye tracking system (SR Research) at a sampling rate of 1000 Hz. All procedures abided by regulations established by the Canadian Council on Animal Care and were approved by the Institutional Animal Care Committee of the Montreal Neurological Institute.

We performed simultaneous recordings in the annectant gyrus, where area V3A is located (Essen and Zeki, 1978; Galletti and Battaglini, 1989), and in the posterior side of the superior temporal sulcus, where area MT is located (Dubner and Zeki, 1971; Gattass and Gross, 1981). To do so, we approached the two areas posteriorly by positioning 32-channel linear microelectrode arrays (Plexon V-Probes with 0.15 mm inter-channel spacing and impedance range between 1-2 MΩ.) so that the uppermost ten channels (spanning ∼1.5mm) were located within area V3A, while the deepest ten channels made contact with ∼1.5mm of area MT. The areas were identified based on anatomical MRIs, depth from the cortical surface, and the transitions between white matter and gray matter along the electrode trajectory. In particular, the presence of a white matter tract makes the separation between V3A and MT clear, as single-unit activity is absent at sites within this region, as shown in Figure 1A.

**Figure 1.**
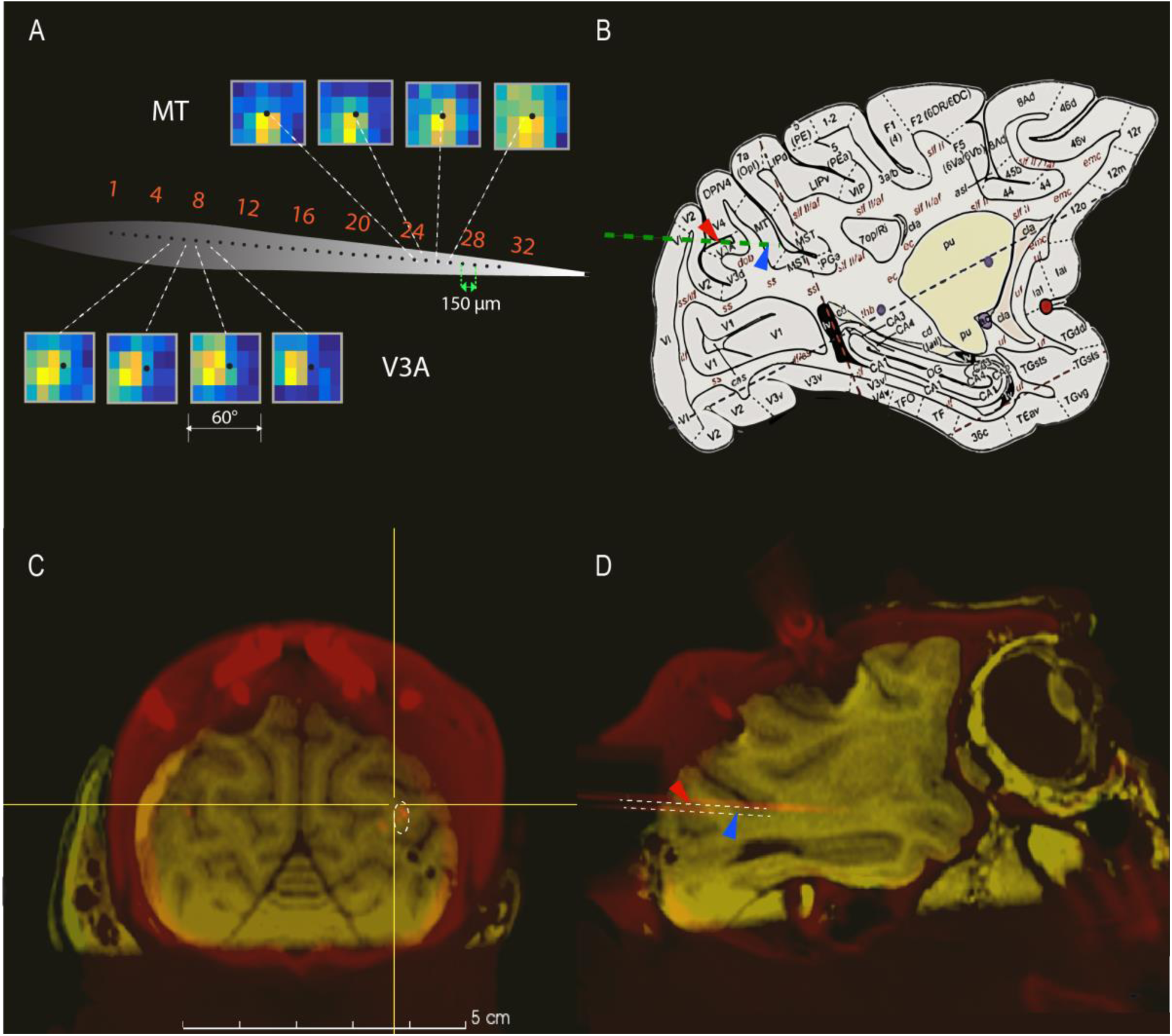
Electrophysiological recording locations and progression of receptive fields across the recording channels. A) Receptive field heat maps for neurons recorded by consecutive sites on the linear probe (indicated by green line in B). Neurons mapped from channels 5 to 8 are located in V3A, while those on channels 24 to 27 are located in MT. Yellow and bright orange indicate stimulus locations that produced the highest firing rates, while blue indicates that the neuron did not fire above baseline at those locations. B) Location of V3A in a standard macaque monkey brain atlas (Saleem and Logothetis, 2012), with the electrode trajectory indicated by a green line. Red arrows in B and D indicate V3A, while blue arrows denote MT. C)& D) Coronal and sagittal view of the MRI from one monkey. The orange dots and lines are the tracks of the recording probe from a CT scan done on the same monkey and registered with the MRI. Dotted white lines are placed to highlight the locations of the recording probe.

The electrode’s position was confirmed with CT imaging. We lowered a stainless-steel guide tube into the recording chamber, positioned a 250 μm tungsten electrode 5mm above the typical recording location, and acquired a single volume with 300 μm isotropic voxels using a Vimago CT scanner (80 kVp). These data were then registered with the MRI, allowing us to precisely visualize the electrode array’s location (Figure 1C, D).

Because we were recording simultaneously from V3A and MT, the vast majority of our data came from one or two electrode trajectories that provided access to both areas in each animal. Recording sites were screened for the presence of clear, visually-driven neural activity during the initial manual mapping, but not for stimulus selectivity. Spike sorting was done by thresholding and filtering the neural signal online, then assigning spikes to single units by a template-matching algorithm (Plexon MAP System). Following this procedure, manual spike sorting was done using a combination of automated template matching, visual inspection of waveforms, clustering in the space defined by the principal components, and absence of absolute refractory period (1 ms) violations (Plexon Offline Sorter). On average, we recorded from 1.42 well-isolated units per electrode channel.

### Visual stimuli

Visual motion stimuli were displayed on an LCD screen with a frame rate of 75 Hz and a resolution of 1280 x 800 pixels; the viewing area covered 60° horizontally and 40° vertically. Animals were trained to fix their gaze at a small dot at the center of the screen. After 400 ms of fixation, a stimulus was shown for another 400 ms. Animals were required to maintain fixation throughout the stimulus presentation to receive a liquid reward. On all trials, gaze was required to remain within 1.5° of the fixation point in order for the reward to be dispensed. Data from trials with broken fixation were discarded.

#### Receptive field mapping

Spatial receptive fields were initially estimated by hand-mapping the population receptive field positions in each area. For a subset of these recordings (77 V3A neurons and 58 MT neurons), we obtained quantitative estimates of receptive field position and size by randomly presenting moving dot patches at random positions within a 4×6 grid. The radius of each patch was 5°, and each position was separated from the neighboring position by 10°. Four different motion directions were incorporated to cover the tuning of the different cortical populations recorded simultaneously. The eccentricities and the sizes of receptive fields were estimated using 2D Gaussian fits to the firing rates of each neuron. We found that receptive fields were centered throughout the contralateral visual space, including the upper and lower visual fields, with eccentricities ranging from 5 to about 40 deg. RFs were generally centered in the contralateral visual field, but we did observe cells with ipsilateral responses as well: 12% (9/77) of V3A cells and 38% (22/58) of MT cells; t-test corrected for multiple comparisons, as reported previously for MT (Raiguel et al., 1995; Raiguel et al., 1997). Fitted RF sizes for large eccentricities (>5°) in MT were slightly larger than those in V3A (p = 0.02, t-test) (Figure 2).

**Figure 2.**
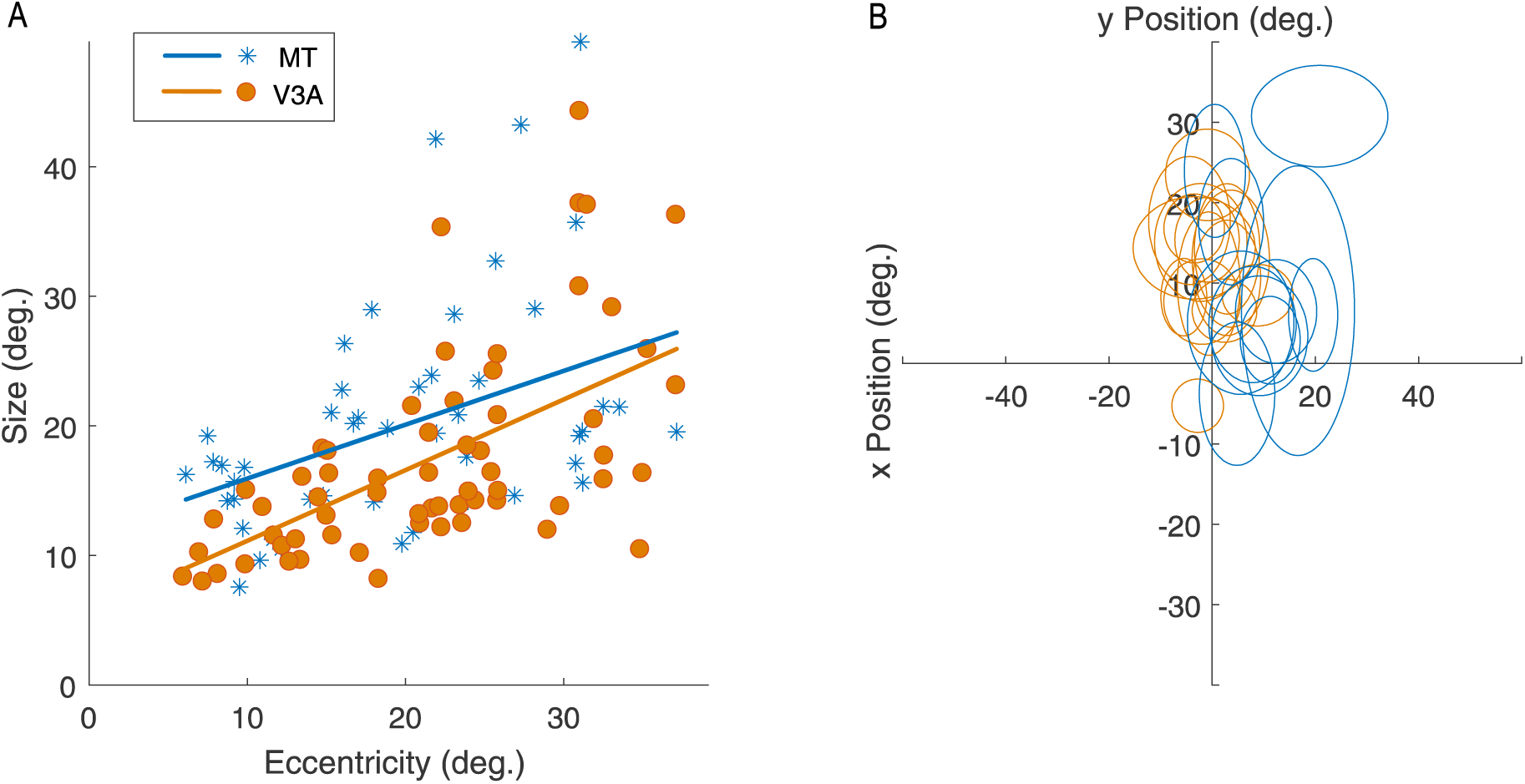
Receptive field sizes and eccentricities in V3A and MT. A) MT cells (blue asterisks) have on average larger receptive fields than V3A cells (orange circles) for the same eccentricities. Linear fits are shown for both populations; orange for V3A and blue for MT. B) Receptive fields from an example recording session. Relationship between RF eccentricity and size with tuning properties are shown in extended figures 2-1 and 2-2.

#### Coherence and Contrast Sensitivity

Based on the mapping of spatial responses at the start of each recording session, the optimal stimulus position was determined for each area. In most sessions, the center of mass of the population response was sufficiently different between V3A and MT that we performed measurements of sensitivity at two different positions, one optimized for each area. In recording sessions in which there were large variations of sizes and/or eccentricities within either area, we used a large stimulus to cover the entire contralateral field.

Measurements of sensitivity were obtained with random dot kinematograms and drifting sinusoidal gratings. Gratings with a spatial frequency of 1 cycle/° were displayed on a gray background (luminance of 98.8 cd/ m^2^). At 100% contrast, the luminance of the sinusoidal grating patch ranged from 0.4 cd/m^2^ to 198 cd/m^2^. Gratings were windowed in a Gaussian envelope, yielding grating patches with radii ranging from 4 to 15°; the radius was adjusted in each recording session according to both V3A and MT receptive field sizes. The stimulus strength and type were chosen randomly on each trial by varying the contrast or coherence (see Results). Because we recorded from V3A and MT simultaneously, the set of coherence/contrast levels was interleaved with two stimulus positions, two stimulus sizes, and eight directions of motion within a block of 10 repetitions per combination. This task was repeated twice to cover two speeds (8 and 16 °/s).

#### Tuning for translation and optic flow motion

The direction tuning of single units was assessed with 100% coherently moving white dot patches on a black background. Dot diameter was 0.1°, and dot density was 2 dots/^(^°^)2^. Each dot had a lifetime of 10 frames, after which it was assigned to a random position inside the patch. On each trial, a single dot pattern appeared, centered at a random locations chosen from a 3 by 3 grid of possible stimulus positions (Figure 3). The 3 by 3 grid was centered around the fixation point, which was at the center of the screen. The patch radius was 10°, and the separation between grid positions was 17° in both the vertical and horizontal directions. This stimulus size elicited robust responses from most neurons, though it unavoidably engaged the inhibitory surrounds as well (Tsui and Pack, 2011; Liu et al., 2018). Although inhibitory surrounds can affect population coding of optic flow (Cui et al., 2013), they do not appear to change single-neuron selectivity dramatically in most MT neurons (Tanaka et al., 1986; Pack et al., 2005; Perge et al., 2005).

**Figure 3.**
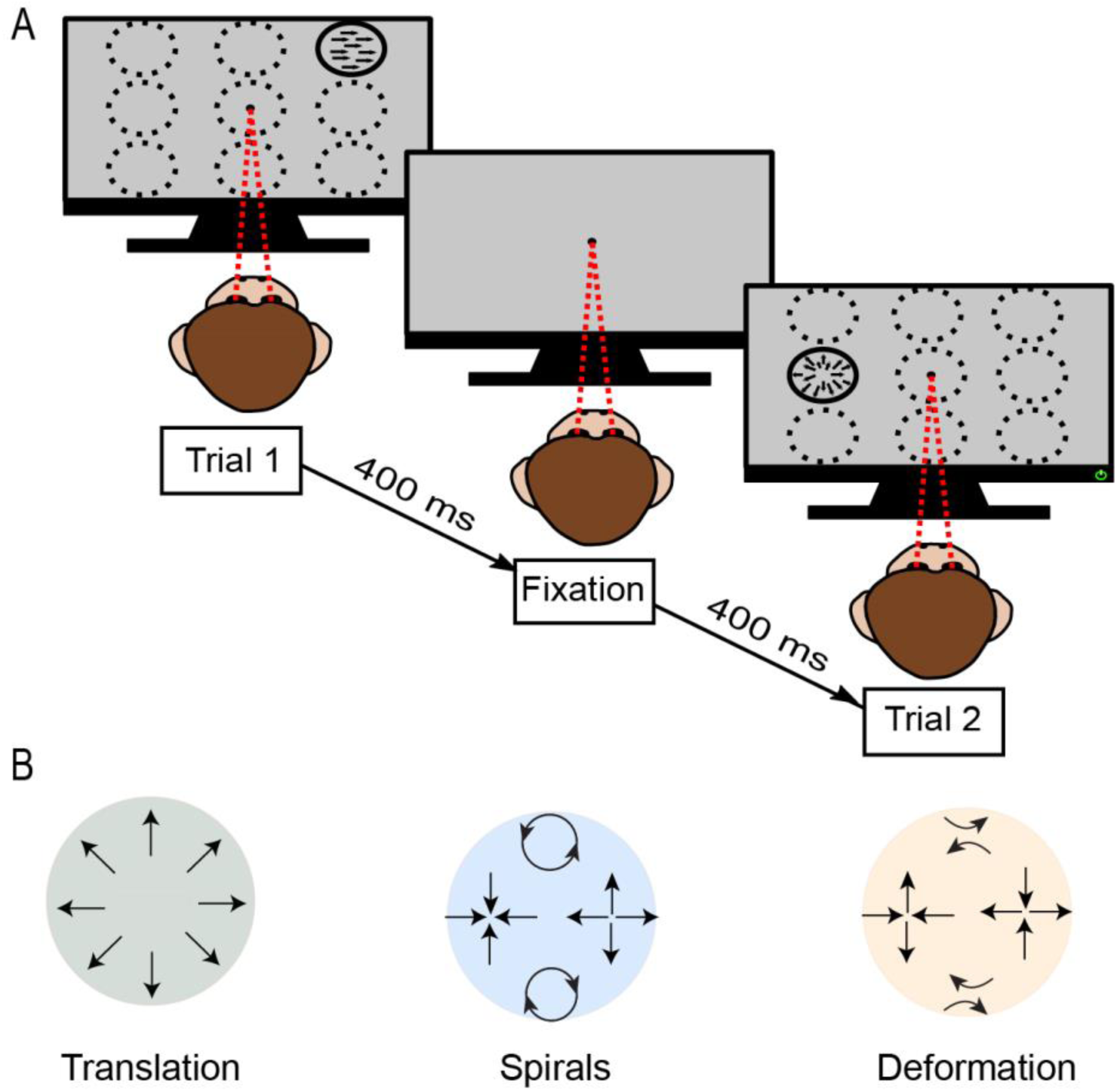
RDK stimuli used in the study. A) Visual stimuli presented to the monkey during recording sessions. The central dot indicates the fixation point at which the monkey maintained gaze during stimulus presentation. The dashed white circles show the 9 possible locations on the screen where the stimulus could be presented on each trial. B) Geometric representations of the different directions for each motion type displayed around a circle.

Tuning was measured over the 3 different motion types (translation, spirals, deformation) defined in equations (1), (2), and (3). Eight directions, represented by θ in the equations below, were defined for each category, sampled at 45° intervals. In translation motion, dots moved in a single direction. The 8 directions in the spiral stimuli corresponded to expansion, contraction, rotation, and their intermediates. Deformation motion included expansion on one axis and contraction on the perpendicular axis, with the axis changing across motion direction. Together, these stimuli span the space of first-order optic flow (Orban et al., 1992).

For translation stimuli, the speed (*v*_*0*_ in equation (1)), ranged from 8 °/s to 32 °/s, though many recordings were done at 16 °/s only. For optic flow stimuli, the speed (*w*_*0*_ in equations (2) and (3)) was set to a value between 3 Hz and 5 Hz. Optic flow speeds were chosen based on an initial, qualitative screening of neuronal responses. The combinations of all the different parameters resulted in 216 different stimuli that were randomly interleaved and repeated 4 or 5 times each.

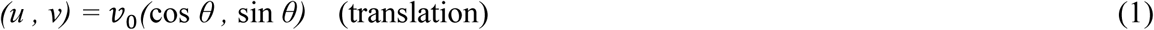

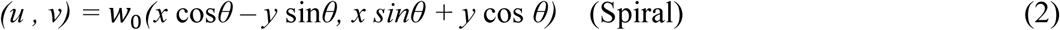

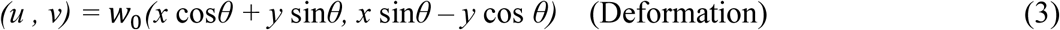

In each of these equations, (*u, v*) represents the instantaneous velocity of a dot at position (*x, y*).

## Data Analysis

Visually driven firing rates were calculated as the mean spike rate during the period from 50 to 400 ms after the onset of each stimulus, averaged across repetitions of the same stimulus. Baseline firing rate was calculated as the mean firing rate during the time from 0 to 400 ms after the onset of fixation, averaged across all trials within a task. Only neurons that responded to visual stimulation with a firing rate increase of at least 2 standard deviations above the mean of the spontaneous firing were included in further analysis. For the neurons that met this criterion, we calculated a Directional Selectivity Index (DSI), defined as 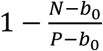 where *P* and *N* are the mean firing rates in the preferred direction and null directions, respectively, and *b*_0_ is the mean spontaneous (“baseline”) firing rate. Mean baseline firing rates were 3.6 ±3.6 Hz for MT and 2.4 ±3.6 Hz for V3A.

To further characterize direction tuning, we calculated circular variance according to the formula: *V* = 1 − |*R*|, where *R* is the mean resultant vector obtained by: 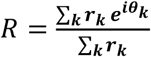

Here *r*_*k*_ is the firing rate for direction *θ*_*k*_ (Fisher, 1995; Jammalamadaka and Sengupta, 2001). A circular variance of 1 indicates that the mean resultant vector for all the directions is zero, and the neuron has no direction preference. A circular variance of 0 means that the neuron responds only to one direction.

For the estimates of response latency, post-stimulus time histograms (PSTHs) were created by binning the spike trains into 15 ms sliding bins. Response latency was calculated as the first point at which the PSTH exceeded the 95^th^ percentile of the spontaneous activity recorded 400 ms before stimulus onset, and which was followed by 2 successive bins that exceeded the 99^th^ percentile of the spontaneous activity (Maunsell and Gibson, 1992). Latency was measured for using the stimulus parameters (direction, speed, and location) that elicited the highest firing rate for each individual neuron.

In order to perform the clustering analysis, multi-unit (MU) activity was extracted from each of the 32 contact sites of the linear probe by including all the waveforms that exceeded a threshold that ranged from −31 to −28 microvolts, depending on the experimental conditions. In order to exclude the correlational effect of single units (SU) on their corresponding MU, we removed every MU spike that occurred within a 1 ms window around each SU spike on the corresponding channel (Shao, DeAngelis et al. 2018). For each channel, we then computed the preferred direction for each SU and MU pair, as the circular mean of firing rates across directions. To the extent that there is clustering for direction selectivity, the SU and MU preferences should be similar at each site. We quantified this similarity by computing the Pearson correlation coefficients between SU and MU preferences across channels.

To estimate the effects of eye position on firing rate, we regressed the mean eye position and firing rates for each neuron, using the equation below.

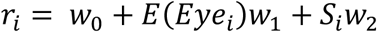

This equation relates the firing rate *r*_*i*_ for each neuron during trial *i* to the stimulus *S*_*i*_ on that trial and the mean of the eye position *Eye*_*i*_, where 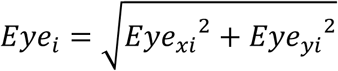. The values of *w*_0_,*w*_1_,*w*_2_ represent the bias term and the weights for eye position and stimulus identity.

### Quantification and Statistical Analysis

All statistical analyses were performed using built-in MATLAB functions and/or custom scripts. *T*-tests were used to test normally distributed variables, such as receptive field sizes. Elsewhere we used the non-parametric Wilcoxon rank-sum test for distributions of circular variances and DSIs that showed non-normal distributions. For contrast and coherence sensitivity analysis, two-sample Kolmogorov-Smirnov tests were used to test the significance of the differences between cumulative distribution functions for the DSIs under each condition in each cortical area. Significance was defined as p < 0.05; all tests were two-tailed.

## Results

Although there is some evidence that neural activity in V3A is concerned with visual motion (Galletti et al., 1990; Tootell et al., 1997; Vanduffel et al., 2002; Anderson and Martin, 2005; Chen et al., 2016), very little is known about the selectivity of single neurons in this area. We therefore recorded from a large population of V3A neurons and characterized their responses to motion stimuli of varying complexity. Simultaneous recordings from MT neurons allowed these responses to be directly compared with a classic dorsal stream motion processing area.

### Basic characterization of neurons

#### Recording sites

As described in the Methods section, we identified area V3A on the basis of MRIs taken before the recordings and a CT scan performed with the recording electrode in place. The position of our recording chamber allowed us to record simultaneously from V3A and MT, using 32-channel linear probes. This arrangement is shown in Figure 1D, which illustrates the recording sites in V3A (red arrow), along with those in MT (blue arrow), superimposed on the scan of one monkey. The location of macaque V3A, taken from a standard atlas (Saleem and Logothetis, 2012), is shown for comparison in Figure 1B. Figure 1A shows a typical progression of receptive fields along the length of the probe from an example recording. In total, we obtained measurements of visual responses in 285 units in area V3A and 231 units in area MT, though not all neurons were tested with all stimuli.

#### Response Latencies

We first analyzed response latencies to visual stimuli with simultaneous recordings from V3A and MT. Response latencies are believed to increase in cortical areas that are higher in the hierarchy of sensory information processing (Raiguel et al., 1989; Vogels and Orban, 1994; Nowak et al., 1995; Schmolesky et al., 1998). We measured the response latency of the simultaneously recorded V3A and MT neurons to moving random dot kinematogram (RDK) stimuli, selecting for each neuron the optimal stimulus conditions (see Methods). For V3A, the response latency ranged from 40 to 243 ms, with a median of 78 ms. Latencies in MT ranged between 40 and 195 ms, with a median of 74 ms, similar to what has been reported previously (Raiguel et al., 1989; Raiguel et al., 1999; Oram, 2010). Though latencies in the two areas were quite similar, a trend toward longer latencies was observed in V3A responses (p = 0.056; Wilcoxon rank-sum test) (Figure 4).

**Figure 4.**
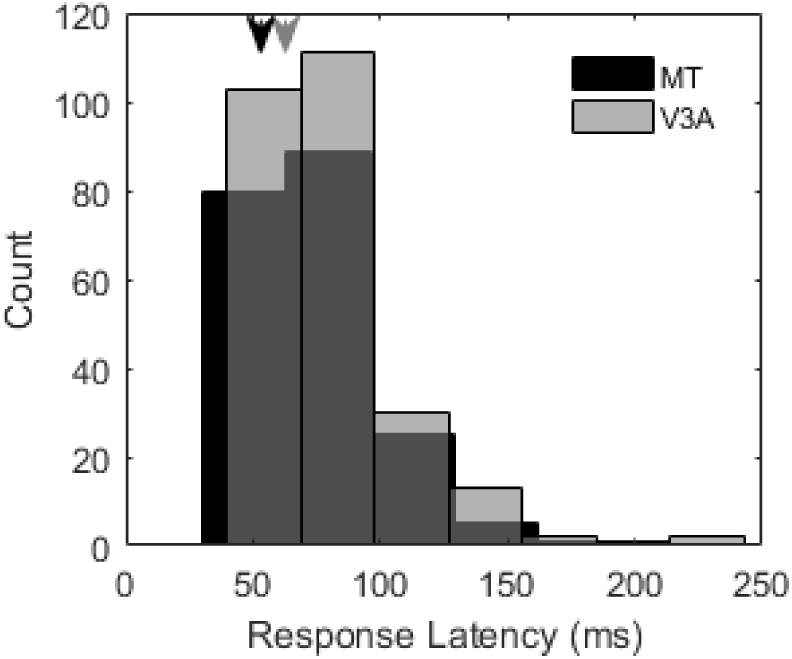
Response latencies for MT and V3A. The distributions of latencies are similar between areas, though the median in slightly highger in V3A. Median values are indicated by arrows on top of the histograms; black for MT and grey for V3A. This figure has two extended figures representing data from each animal separately.

### Selectivity for translation motion

#### Neural selectivity for gratings and RDKs

We first assessed direction selectivity in the V3A population using 100% coherent RDKs undergoing translation motion. When tested in this way, 217 out of 276 cells (79%) exhibited high direction selectivity, defined here as a direction selectivity index (DSI) of at least 0.5. Similarly, when tested with drifting gratings, 188/242 of V3A neurons (78%) exhibited high direction selectivity. Across the population, the median DSI, computed from responses to coherent RDKs, was 0.74.

By comparison, in MT we found that 82% (181/221) of the neurons exhibited high direction selectivity for RDKs and 80% (165/205) for gratings, with the median DSI being 0.77. Note that this level of direction selectivity is similar to early reports of recordings in the posterior bank of the superior temporal sulcus (STS) where MT is located (Dubner and Zeki, 1971); however it is somewhat lower than reported subsequently for MT (Albright, 1984; Britten et al., 1992) in experiments that optimized the stimulus for each neuron. We attribute this difference to our use of multisite recordings, wherein the stimulus cannot be optimized per cell, and indeed similar levels of direction selectivity have been reported in MT for non-optimal stimulus conditions (Raiguel et al., 1995). Across both classes of stimuli, the DSI distribution for MT neurons was not significantly different from that for V3A neurons (p = 0.50; Wilcoxon rank-sum test).

In V3A, 80.3% (195/243) of the tuning curves were well fit by a von Mises (circular Gaussian) functions (r^2^ > 0.5). The figure for MT was similar (143/177 or 80.8%). The median tuning widths for these cells were 83^°^ in V3A and 91^°^ in MT, with the latter figure being similar to what has been reported previously in MT (Albright, 1984; Britten and Newsome, 1998). Thus although previous work has not identified a clear specialization for motion in V3A neurons (Gaska et al., 1988), our results suggest that the representation of direction in this area is comparable to that found in a classic motion processing dorsal stream region.

Of the two animals used in the study, one had undergone extensive training in a motion psychophysics task (Liu and Pack, 2017), while the other had never been trained for any behavior other than fixation. Nevertheless, the distributions of DSI values were not significantly different between the two animals (p = 0.20; Wilcoxon rank-sum test). In the following sections, we combine data across animals, but a summary of the results in each animal can be found in the extended data (Figures 7-1, 7-2, 4-1 and 4-2).

#### Sensitivity to visual motion stimuli

We next examined the sensitivity of direction selectivity in V3A and MT to stimulus strength, using modulations of contrast and motion coherence. Because we were interested in the representation of motion direction, we discarded cells that had a DSI below 0.3 for this analysis. This left 224 of 252 cells in V3A and 183 of 196 cells in MT. All cells were tested at two levels of stimulus contrast (for gratings) or motion coherence (for RDKs).

Figure 5 shows the cumulative DSI for the different stimuli in V3A and MT. No significant differences between areas for the cumulative distributions of DSI were seen at 100% contrast for gratings (p = 0.17, Two-sample Kolmogorov-Smirnov tests) or 100% coherence for RDKs (p = 0.20 for RDKs, Two-sample Kolmogorov-Smirnov tests).

**Figure 5.**
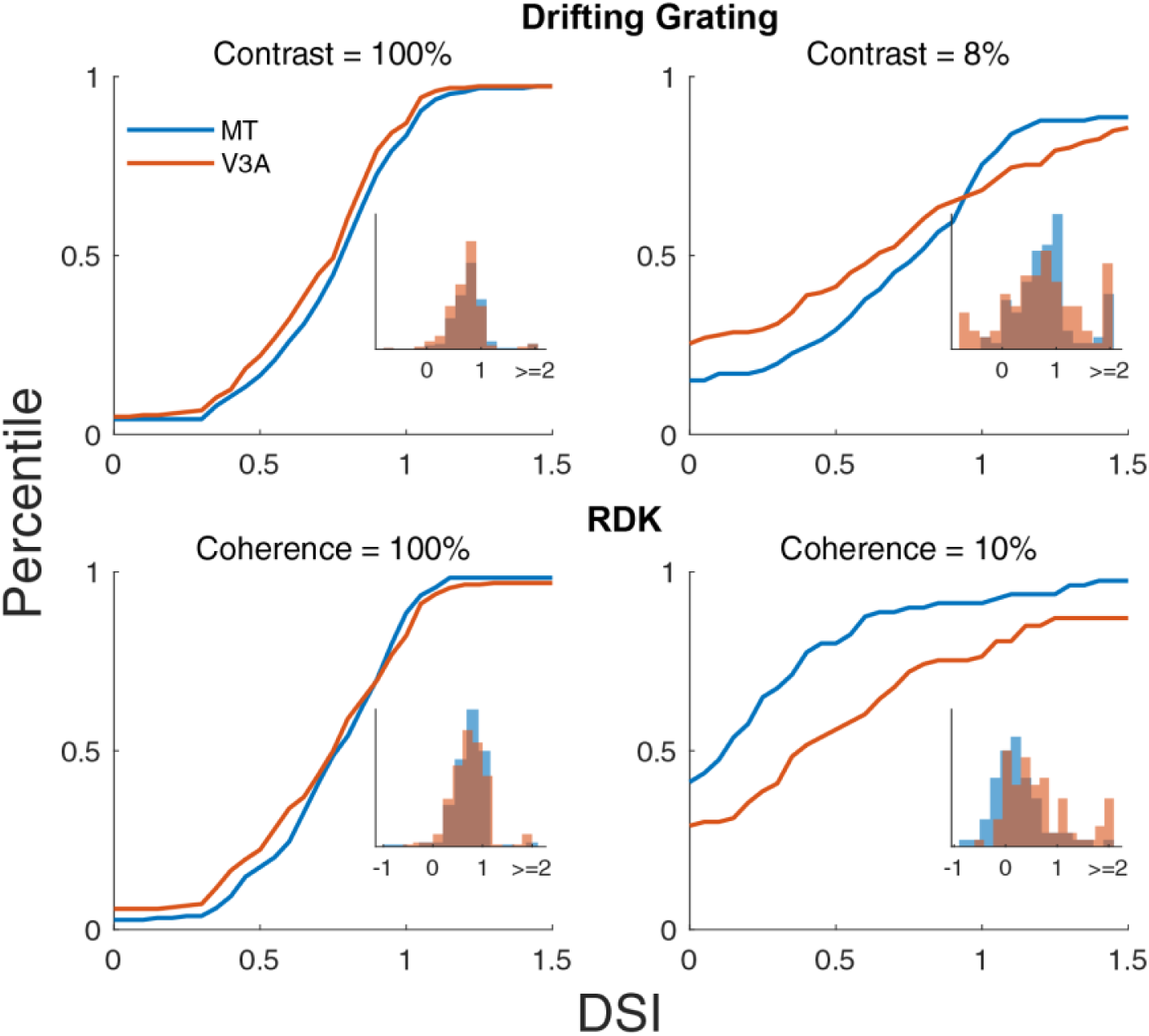
Sensitivity to motion signal strength. Cumulative distribution functions for the DSI in V3A (Orange) and MT (Blue) at different constrasts (for grating stimuli) and coherences (for RDKs). The insets show the distribution of DSIs for each population at each contrast or coherence. Note that negative values can occur when the cells lose direction selectivity, in which case the DSI is driven by random fluctuations in the response.

Cumulative selectivity was also similar between the two areas at low contrast (8%) (p = 0.13, Two-sample Kolmogorov-Smirnov test). Surprisingly, at 10% coherence, cumulative direction selectivity was significantly higher in V3A than in MT (p = 0.001, Two-sample Kolmogorov-Smirnov test). This suggested a possible role for V3A in motion integration, which we consider in the following sections.

### Selectivity for optic flow motion in V3A and in MT

#### Direction selectivity

Previous imaging work has suggested that V3A is involved in representing complex optic flow motion stimuli. These include optic flow components, such as expansion, rotation, and deformation (Koyama et al., 2005; Wall et al., 2008; Furlan and Smith, 2016), which along with translation, span the space of all first-order optic flow (Orban et al., 1992; Mineault et al., 2012). We therefore studied neural responses in V3A (N = 257) and MT (N = 193) to a range of optic flow motion stimuli, including the translation motion described above. All stimuli were comprised of 100% coherent dot patterns (see Methods section for detailed description), presented at nine different locations in the visual field (Figure 3).

Figure 6 shows the responses of two example V3A cells to our battery of motion stimuli.

**Figure 6.**
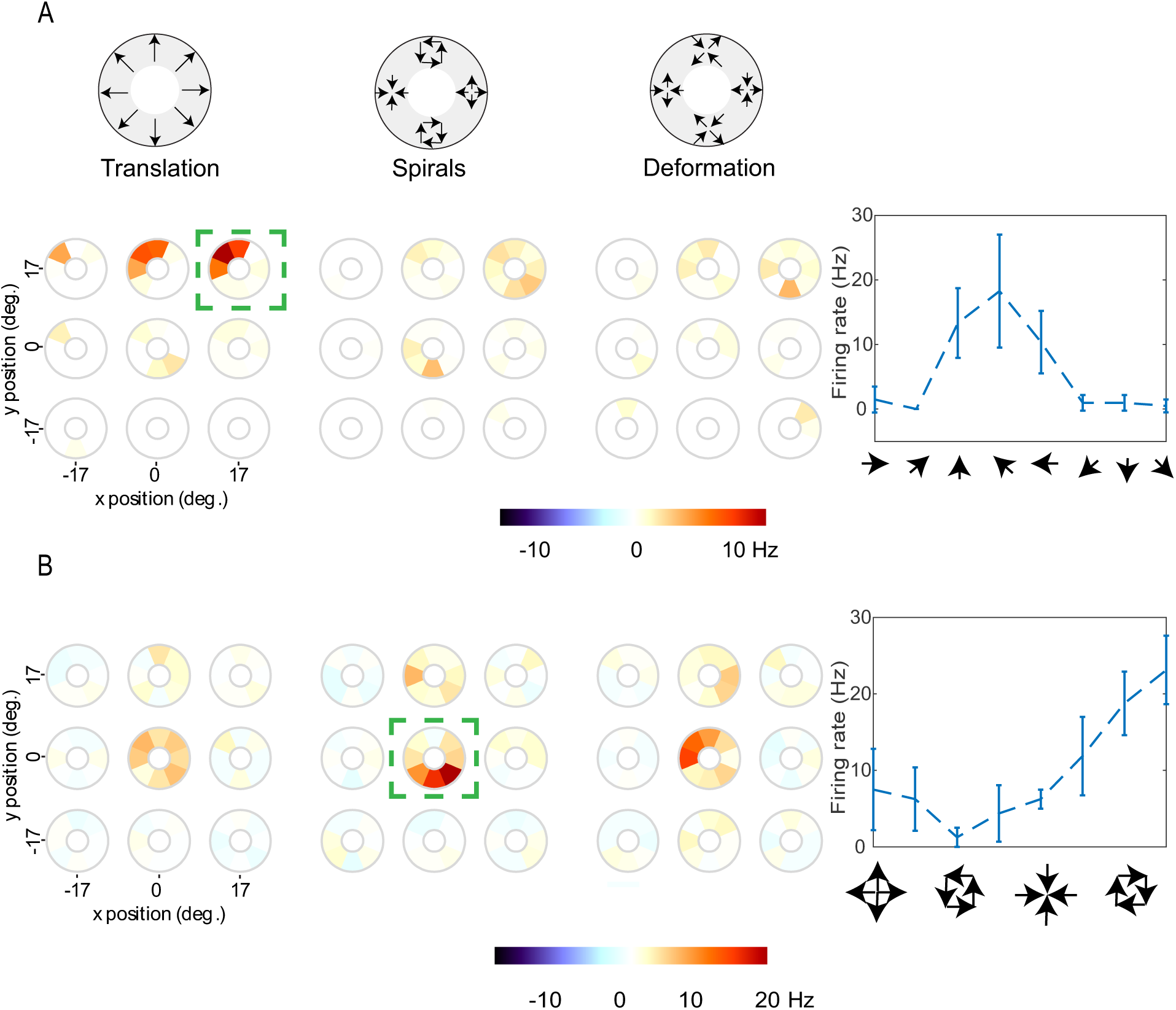
Tuning Curves for two example V3A cells. Within each motion type, there are nine different tuning curves, corresponding to the nine different stimulus positions. At each position, each circle (tuning wheel) represents the firing of the neuron for each of 8 motion directions; orange and red correspond to high firing rates, white to baseline, and blue to firing rates lower than baseline. The green dashed squares indicate the position and the motion type of the stimulus that elicited highest firing. The tuning curve inside the green square is shown in Cartisian coordinates to the right of the tuning wheels. A) Tuning curves for a translation selective V3A cell. The color bar indicates the range of firing rates in Hz. B) Same as A but for another neuron that is selective for spirals. Average tuning curves for MT and V3A are shown in figure 6-1 in the extended data.

**Figure 7.**
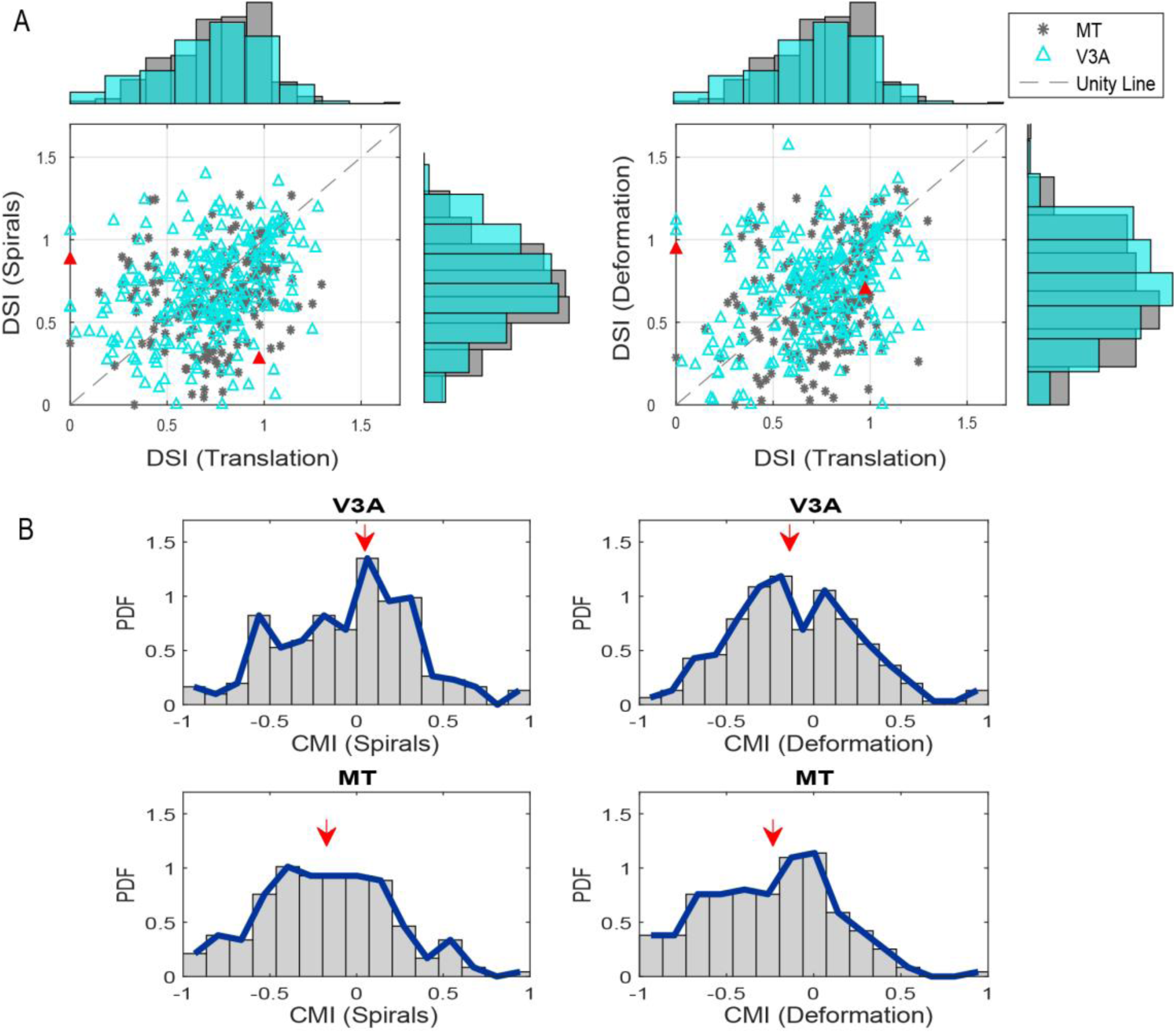
V3A responses to optic flow stimuli. A) Scatter plots for V3A DSIs (cyan) and MT DSIs (grey) for spirals versus translation (left) and deformation versus translation (right). The unity line is shown by a dashed gray diagonal line. The red triangles refer to the example neurons shown in Figure 6. B) Probability density function of the complex motion index (CMI) for sprials (left) and and for deformation (right) for V3A (upper panels) and MT (lower panels). A value of zero for each CMI indicates no preference for complex optic flow over translation motion, while positive numbers indicate a preference for complex optic flow and negative numbers a preference for translation. The red arrows indicate the median for each distribution. This figure has two extended figures representing data from each animal separately.

For each cell, we show the responses to three types of motion: translation, spiral, and deformation (explained in the Methods section). Within each motion type, there are nine different tuning curves, corresponding to the nine different stimulus positions. At each position, each circle represents the firing of the neuron for each of 8 motion directions; orange and red correspond to high firing rates, white to baseline, and blue to firing rates lower than baseline (Figure 6). The example V3A cell in Figure 6a is highly selective for translation motion and specifically for motion around 135°, while its responses to other types and directions of motion are weaker. This cell therefore exhibits direction selectivity for translation motion at a particular point in visual space. In contrast, the example V3A cell in Figure 6b exhibits selectivity for optic flow motion: It responds weakly to translation motion (left) but has a strong response to clockwise rotation in the center of the stimulus array (middle). It also responds well to deformation with a specific axis of motion (right).

To quantify the selectivity for stimulus motion across the V3A and MT populations, we used the DSI to represent neural selectivity for translation, spirals and deformation in each area. For each neuron, direction selectivity was assessed for each motion type at the optimal stimulus position, as determined from data of the type shown in Figure 6.

Across the neural population, we found a wide variety of tuning preferences in both areas. Nevertheless, pairwise comparisons between areas indicated that DSIs for spiral and deformation motion were higher in V3A than in MT (spiral: p = 0.03; deformation: p = 0.09; Wilcoxon rank-sum tests), while there was no significant difference between areas for translation motion (p = 0.5, Wilcoxon rank-sum test).

Of course, the observed selectivity for optic flow motion stimuli could simply be a consequence of selectivity for translation motion. This would occur if the neurons were responding to local motion within each stimulus (Lagae et al., 1994), rather than to the motion pattern *per se*. In this case, the DSIs for spirals and deformation should be correlated with the DSIs for translation. This was indeed the case in MT (Pearson’s r = 0.22, p = 0.003 for translation and spiral DSIs, r = 0.31, p ≪ 0.01 for translation and deformation DSIs) and in V3A (Figure 7a; Pearson’s r = 0.33, p ≪ 0.01 for translation and spiral DSIs, r = 0.34, p ≪ 0.01 for translation and deformation DSIs). However, the observed correlations were weak, and it is clear from Figure 7A, as well as the example in Figure 6B, that some V3A cells exhibited high DSIs for optic flow motion but low DSIs for translation motion.

To assess the relative selectivity for optic flow and translation motion, we defined a complex motion index (CMI) as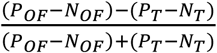, where *P*_*OF*_ and *N*_*OF*_ are the mean firing rates for the preferred and null directions of optic flow motion, with *P*_*T*_ and *N*_*T*_ being defined analogously for translation. We computed the CMI separately for spirals and for deformation.

A negative CMI indicates greater selectivity for translation, while a positive CMI indicates greater selectivity for optic flow.

Figure 6B show the CMI distributions in both areas. In MT, there is a bias toward negative values, with the median CMI being −0.14 for spirals and −0.22 for deformation. In V3A the CMI distributions are shifted towards the positive side of the axis relative to the MT distributions, with the median being 0 for spirals and −0.14 for deformation. Overall, the CMIs for V3A were significantly higher than the CMIs for MT for both spiral and deformation motion (p = 0.001, p = 0.002, Wilcoxon rank-sum test).

#### Tuning width

In addition to response amplitude, a complementary measure of stimulus preferences is the narrowness of tuning curves for different motion types. To quantify this kind of selectivity in the V3A population, we used the circular variance of tuning curves like those shown in Figure 6. Circular variance provides an estimate of the tightness of tuning, and we compared these measures for the three motion types in V3A and MT. As in the previous analysis, we selected the optimal stimulus position from data of the type shown in Figure 6.

The results show that MT neurons typically had a higher circular variance for optic flow motion than for translation motion (p ≪ 0.001, Wilcoxon rank-sum test) (Table 1). V3A, on the other hand, showed a smaller difference between circular variance for optic flow motion and circular variance for translation motion (median 0.78 for spiral, 0.78 for deformation and 0.74 for translation motion; p =0.01, Wilcoxon rank-sum test).

**Table 1.**
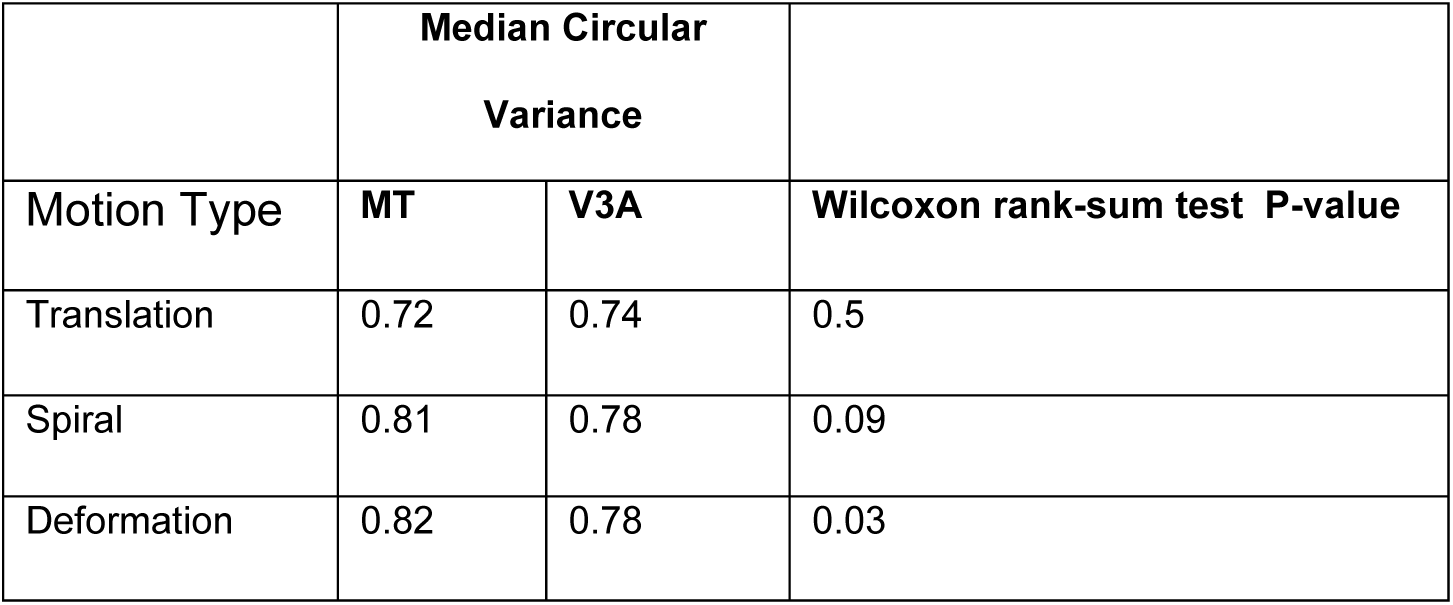
Summary of circular variance in V3A and MT for 3 motion types (translation, spirals, and deformation).

|  | Median Circular<br>Variance |  |  |
| --- | --- | --- | --- |
| Motion Type | MT | V3A | Wilcoxon rank-sum test P-value |
| Translation | 0.72 | 0.74 | 0.5 |
| Spiral | 0.81 | 0.78 | 0.09 |
| Deformation | 0.82 | 0.78 | 0.03 |

Cross-area comparisons revealed that circular variance for translation motion was not significantly different between V3A and MT (p = 0.50, Wilcoxon rank-sum test), while circular variance for optic flow motion showed a tendency to be lower in V3A than in MT. This tendency was significant in the case of deformation stimuli (p = 0.09 for spirals and p = 0.03 for deformation, Wilcoxon rank-sum test). Circular variance was not significantly correlated with RF size or eccentricity (linear regression, p>0.05). Similarly, we found no clear relationship between complex motion selectivity and RF size or eccentricity (Figures 2-1 and 2-2). Together with the results of the previous section, our findings suggest that compared to neurons in MT, neurons in V3A exhibit narrower tuning and somewhat more firing rate modulation for optic flow motion.

#### Clustering

Neurons in most sensory cortical areas exhibit topographical clustering of selectivity (Hubel, Wiesel, & Stryker, 1978; Mountcastle, 1957): Nearby neurons encode similar stimulus features. To examine such clustering in V3A, we calculated the preferred direction of each single unit (SU) and compared it to the selectivity of the simultaneously recorded multi-unit activity (MU). Because the MU represents the summed contribution of many nearby neurons, it is a reasonable proxy for columnar selectivity (Shao et al., 2018). As described in the Methods, we first removed the SU spikes from our estimate of MU, to avoid inflated correlations between the SU and MU preferred directions. These correlations, computed across channels of the linear electrode arrays, served as a metric of clustering for preferred direction (Shao et al., 2018).

For translation we found significant correlations between the SU and MU selectivity for both V3A (t-test, p = 0.0005) and MT (t-test, p = 0.0008). The latter result was expected based on the known clustering properties of MT (Albright, 1984). Clustering in MT was stronger than in V3A for translation (0.51 vs 0.41) and for spirals (0.2, p=0.15 vs 0.019, p=0.88), suggesting a slightly more precise organization in MT than in V3A. In response to deformation, stronger correlations between SU and MU were observed for V3A neurons (0.08, p = 0.45) than for MT neurons (−0.006, p = 0.96), though neither relationship was statistically significant. Thus while both V3A and MT have clustering for translation, we did not find the kind of clustering based on optic flow that has been reported in higher-level areas (Geesaman et al., 1997; Britten, 1998). Since our recordings were limited to a small region within V3A, further experiments are needed to characterize the full topography of this area.

#### Eye movements

Previous work has found that V3A cells receive oculomotor inputs that allow them to distinguish between retinal motion and reafferent motion caused by eye movements (Galletti and Battaglini, 1989; Galletti et al., 1990). Although our experiments involved only fixation, we recorded eye position continuously and were therefore able to assess whether small fixational movements or microsaccades influenced firing rates. To this end, we performed a regression analysis that related distributions of eye positions to firing rate (see methods).

The results showed that the eye position on a trial-by-trial basis did not exert a significant influence on firing rates. The recovered weight of eye signal in the regression model was 0.000005 ± 0.0002, which was not significantly different from zero (t-test, p = 0.6). This analysis, along with previous findings (Bair and O’Keefe, 1998) suggest that small eye movements are unlikely to have affected measures of stimulus selectivity in our experiments.

## Discussion

Using single-unit recordings and a large battery of visual stimuli, we have identified a neuronal population in area V3A that exhibits a high prevalence of direction selectivity. Most of the neurons we encountered were well-tuned for translation motion of drifting gratings or random dots (Figure 6A), and others were well-tuned for more complex optic flow patterns (Figure 6B). The overall profile of neural selectivity in V3A was similar to that observed in a population of simultaneously recorded MT neurons. These results suggest that V3A, or at least some domains therein, should be considered as part of the motion-processing machinery of the primate visual cortex.

### Functional anatomy of V3A

Area V3A has classically been localized as being in the annectant gyrus and the anterior bank of the lunate sulcus (Essen and Zeki, 1978; Zeki, 1978; Galletti and Battaglini, 1989). As a result, it is inaccessible to optical imaging and difficult to pinpoint with electrode recordings. Intersubject variability contributes further to the difficulty in assigning a consistent brain location to the label V3A (Zhu and Vanduffel, 2019). Unsurprisingly, these complications have led to some variability in the functional characteristics reported by different studies.

V3A was initially defined by Van Essen and Zeki in 1978 as a distinct visual area located in the region between V3 and V4 in the lunate and parieto-occipital sulci (Essen and Zeki, 1978), but subsequent studies used somewhat different boundaries and assigned different names to different sub-regions (Felleman and Van Essen, 1987; Colby et al., 1988; Kaas and Morel, 1993). As a result, it is difficult to make direct comparisons between our results and those of previous single-unit papers. For example, based on more recent anatomical boundaries (Saleem and Logothetis, 2012), most of the recording sites in the paper by Gaska et al. (1988) would be classified as V3d, while those in the work of Galletti et al. (1990) would include sites in V3. In our work, we have not attempted to perform anatomical reconstructions of our recording sites, but our *in-vivo* (Figure 1) imaging shows them to be consistently in the dorsal part of V3A, close to the boundary with V4 (Saleem and Logothetis, 2012). Thus while the proportion of direction selective neurons in our study is higher than previously reported (Gaska et al., 1988), it is clear that our recordings also came from a different location.

A related possibility is that V3A (or the V3 complex as a whole) contains patches of cells that are selective for motion direction, interspersed with patches that are selective for other stimulus feature (orientation, color, etc.). This would not be surprising given the evidence for similar functional domains in neighboring areas like V2 (Livingstone and Hubel, 1987; Lu et al., 2010) and V4 (Ghose and Ts’o, 1997; Li et al., 2013). Functional clustering in V3A has been shown previously for gaze modulation of neural firing rates (Galletti and Battaglini, 1989) and for stereoscopic depth encoding (Anzai et al., 2011). In addition, anatomical evidence show that the feedforward projections from V2 to layer 4 of V3A form stripe-like patches, similar to the projections from the thalamus to V1 and from V1 to V2 (Anderson and Martin, 2005). Since our electrode trajectories were constrained by the need to target MT and V3A simultaneously, we have not extensively explored the topography of V3A.

### Stimulus representations in V3A

In addition to the complexity of localizing V3A, there has been considerable debate about its stimulus encoding properties. Imaging studies in humans have found evidence for strong direction selectivity in V3A (Tootell et al., 1997; Smith et al., 1998; Sunaert et al., 1999; Braddick et al., 2000; Seiffert et al., 2003; Koyama et al., 2005; Cardin et al., 2012; Furlan and Smith, 2016; Strong et al., 2017) that is second in strength only to that of the hMT/V5 complex (Tootell et al., 1997; Goebel et al., 1998). At the same time, similar imaging studies in macaques found little direction selectivity in this area (Vanduffel et al., 2001; Vanduffel et al., 2002; Orban et al., 2003). These results have led to the suggestion that V3A has different functional properties in macaques and humans (Vanduffel et al., 2001; Vanduffel et al., 2002; Orban et al., 2003). However, an alternative view, consistent with our results, is that at least some domains within V3A are largely dedicated to motion processing. Because these regions are necessarily smaller in the macaque, they might have been inaccessible to standard fMRI measurements.

A similar discrepancy between fMRI and single-unit findings emerged for the role of V3A in the processing of binocular disparity. fMRI studies in human V3A found that it is more strongly modulated by stimulus disparity than most other visual areas (Mendola et al., 1999; Rutschmann et al., 2000; Backus et al., 2001), and a similar result was found in macaques using functional imaging (Tsao et al., 2003). In contrast, single-unit recordings did not reveal an unusual level of selectivity for disparity (Anzai et al., 2011). Indeed, they found that typical selectivity for disparity in V3A was similar to that in other extrastriate areas.

Finally, there has been some dispute about visual field representations in V3A and related areas. Previous studies have indicated different representations in New World and Old World monkeys (Gattass et al., 1988; Angelucci and Rosa, 2015). Classically V3A represents the central and upper visual field (Essen and Zeki, 1978) while, in New World monkey studies, it contains representations of bother upper and lower contralateral visual fields. However, recent work with higher-resolution imaging suggests that the organization is likely more similar between the species than previously supposed (Zhu and Vanduffel, 2019). Our recordings (in Old World monkeys) covered both the upper and lower contralateral fields and were carried out in the annectant gyrus, which corresponds to the classical location of V3A, as well as the new boundaries revealed in (Zhu and Vanduffel, 2019).

### The position of V3A in the hierarchy of visual motion processing

V3A has classically been positioned at a lower stage of processing than area MT (Felleman and Van Essen, 1991; Felleman et al., 1997), and human imaging studies have provided some support for this idea (Huk and Heeger, 2002). However, anatomical projections between MT and V3A are of the intermediate type, meaning that neither one of the areas projects exclusively to input layers of the other; rather the projections are spread evenly across all cortical layers (Ungerleider and Desimone, 1986). Some of these projections extend contralaterally, a pattern that is also seen in MT’s projections to MST and VIP (Ungerleider and Desimone, 1986).

From a physiological standpoint, our results indicate that motion processing is generally similar between V3A and MT (Figures 5 and 7). Thus, at the level of single-neuron stimulus representations, there is little reason to suggest that V3A is at an earlier stage than MT, and indeed, there is some evidence that V3A represents complex optic flow stimuli more effectively than MT (Figures 7B). Moreover, the response latencies of V3A neurons are on average slightly higher than those in MT (Figure 4), and V3A is more sensitive to low-coherence motion stimuli than MT (Figure 5).

The latter result might seem surprising in light of the fact that receptive fields in MT are somewhat larger than those in V3A (Figure 2). One way to overcome noise in low-coherence RDKs is to integrate information over space, and so one might expect MT neurons with larger receptive fields to exhibit greater sensitivity to such stimuli. However, psychophysical results suggest that this is not the case, as motion direction in larger RDK stimuli is not necessarily perceived more accurately (Bakhtiari et al., 2020). Instead, sensitivity to low-coherence RDKs might be related to the integration of inputs with different preferences for direction or spatiotemporal frequency (Smith et al., 1994; Bakhtiari et al., 2020). If V3A neurons integrate more widely in the direction domain, it might also account for the enhanced selectivity for optic flow mentioned above (Khawaja et al., 2013).

A related point is the existence of ‘real motion’ cells in V3A (Galletti et al., 1990) that respond to external motion when the eye is stationary, but not to similar retinal motion caused by rotation of the eye. These results suggest that an input from oculomotor regions reaches area V3A (Galletti et al., 1990), and a similar conclusion has been reached in human studies (Fischer et al., 2012). By comparison, cells of this type were not often found in MT (Newsome et al., 1988; Erickson and Thier, 1991; Chukoskie and Movshon, 2009), but they were found in high-level area MST (Newsome et al., 1988; Tanaka and Saito, 1989; Shenoy et al., 1999; Britten and van Wezel, 2002; Inaba et al., 2007). These results, combined with the above-mentioned optic flow motion selectivity, suggest a possible role for V3A in self-motion perception. Indeed, disrupting V3A with TMS impairs the ability to localize heading direction (Strong et al., 2017).

These considerations lead to the possibility that V3A is more similar to neighboring area V6 (Galletti et al., 1995) than to lower-level visual areas like V1 or V2. Like V3A, V6 is located in the lunate sulcus, although it is situated far more medial than our recording sites. Neurons in V6 also exhibit sensitivity to optic flow stimuli comparable to that found in high-level areas like MST (Fan et al., 2015). A similar selectivity for optic flow was also reported with fMRI in humans (Cardin and Smith, 2010). Non-visual inputs representing eye position were not found in V6 (Fan et al., 2015), but the prevalence of real-motion neurons is higher in this area than in V3A (Galletti and Fattori, 2003). These results suggest a role for both V3A and V6 in orienting or reaching toward moving objects (Galletti and Fattori, 2003).

## Acknowledgements

We thank Julie Coursol, Cathy Hunt, and Dr. Fernando Chaurand for assistance with animal care and preparation. This work was supported by a grant from the Canadian Institutes of Health Research to C.C.P. (MOP-115178).

