## Supplementary figures and images for "Neural selectivity for visual motion in macaque area V3A"

### Figure 2-1

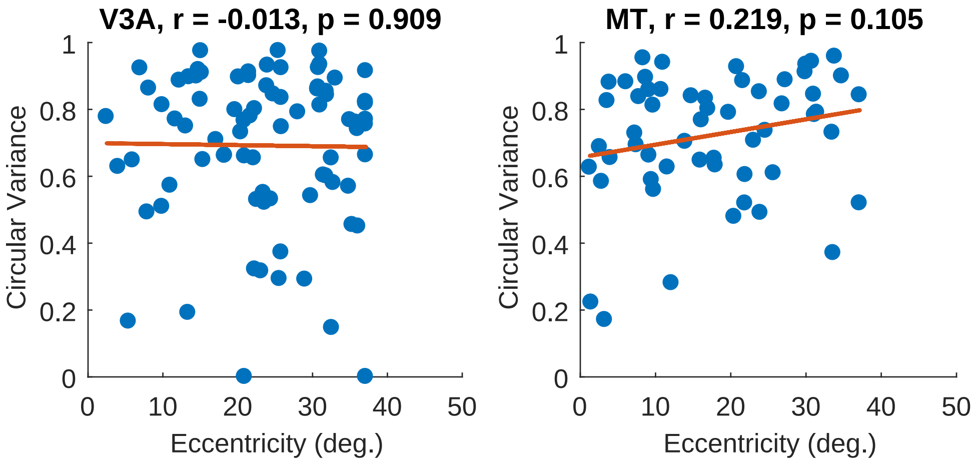

### Figure 2-1

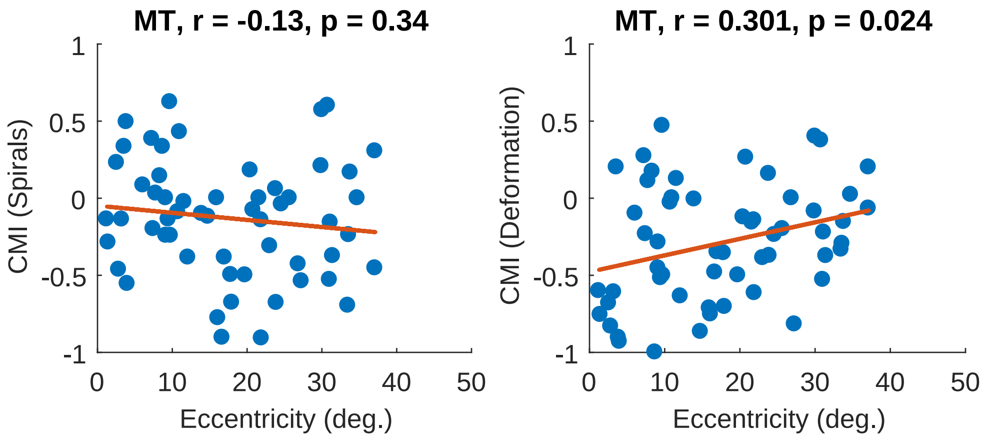

### Figure 2-1

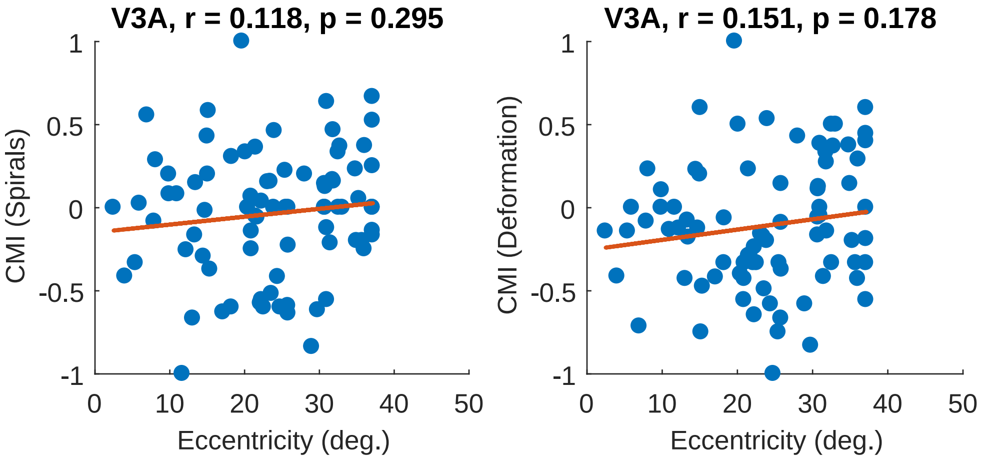

### Figure 2-2

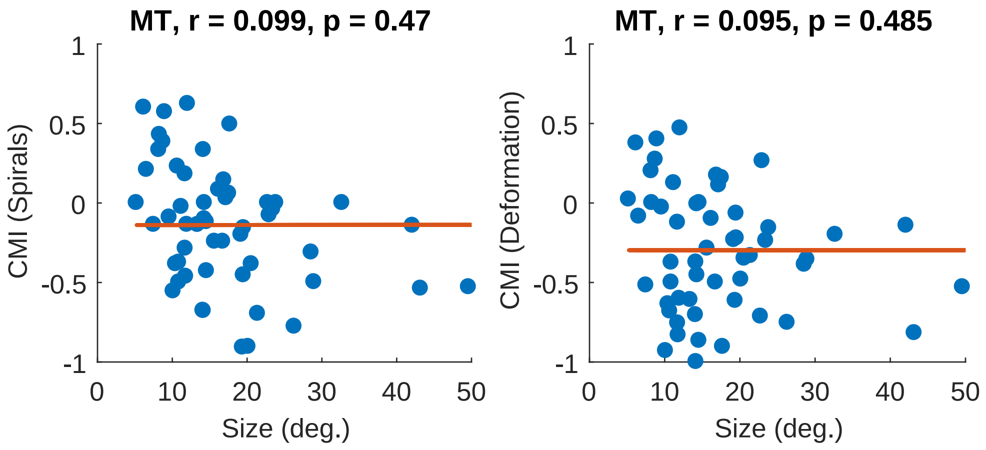

### Figure 2-2

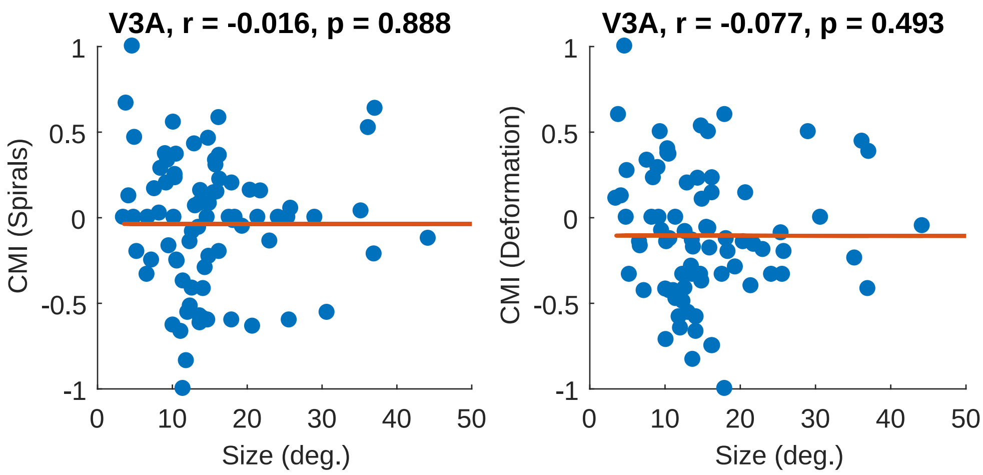

### Figure 2-2

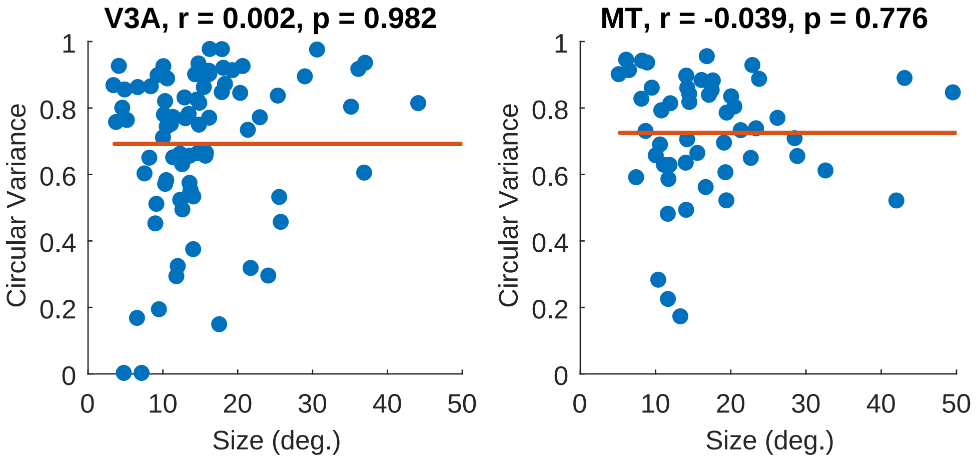

### Figure 4-1

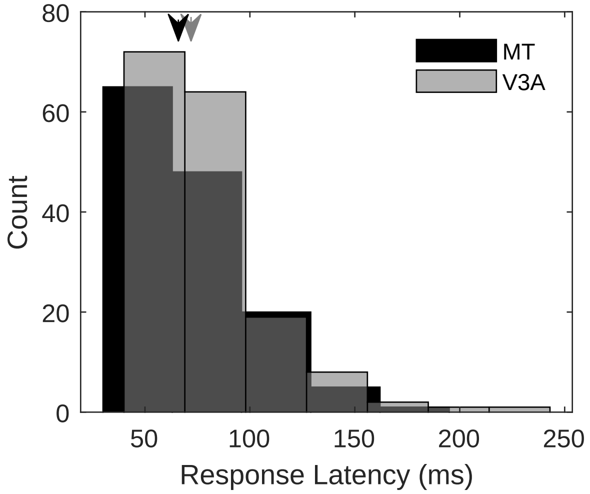

### Figure 4-2

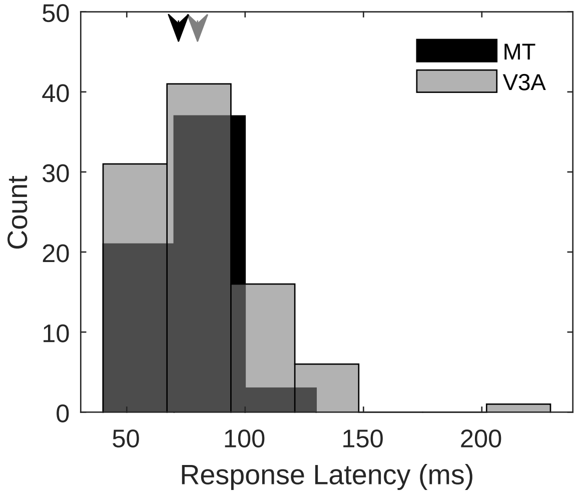

### Figure 6-1

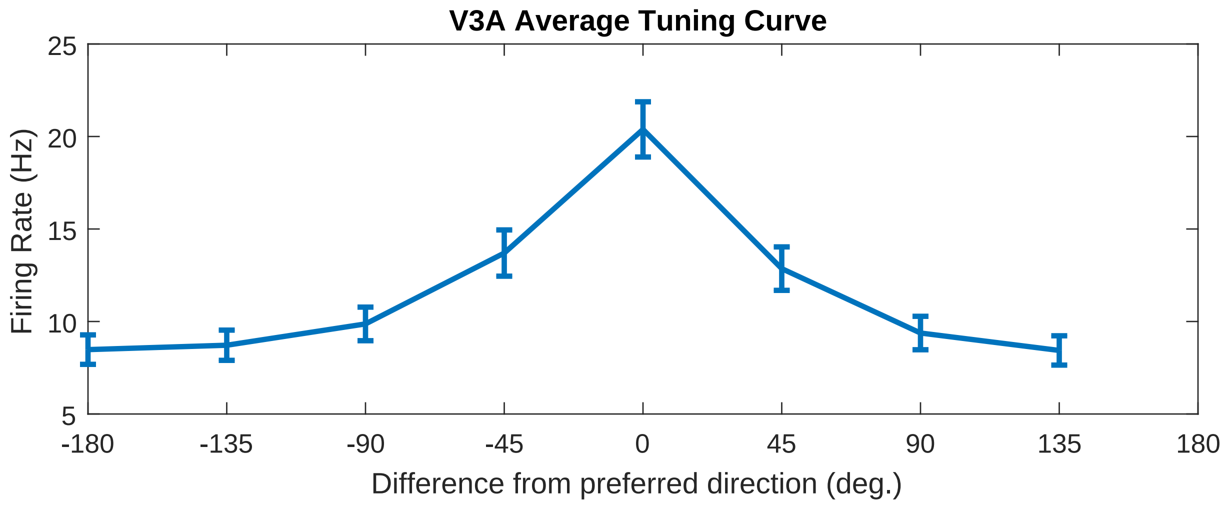

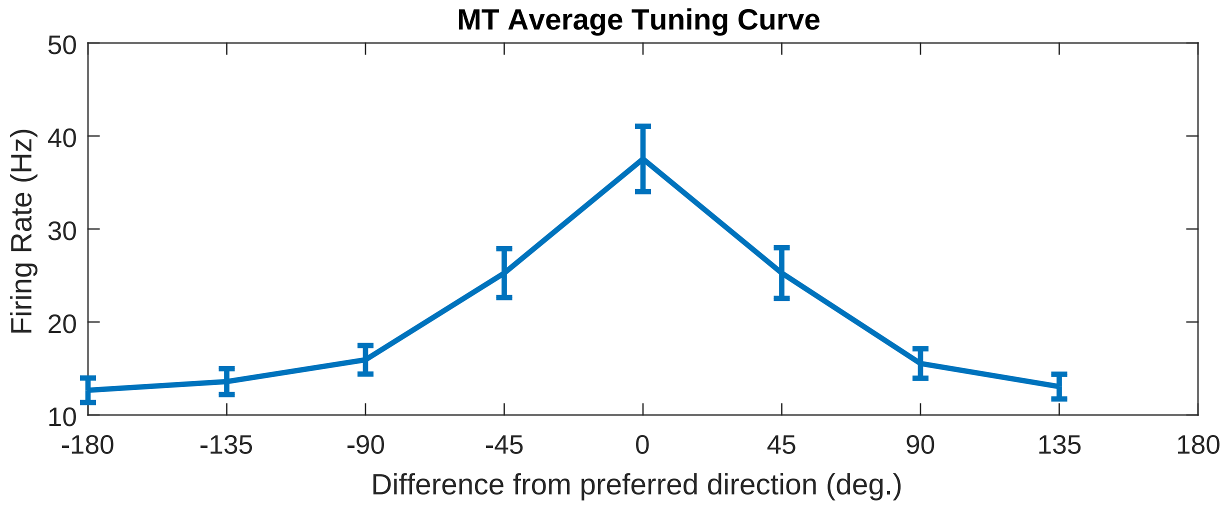

### Figure 7-1

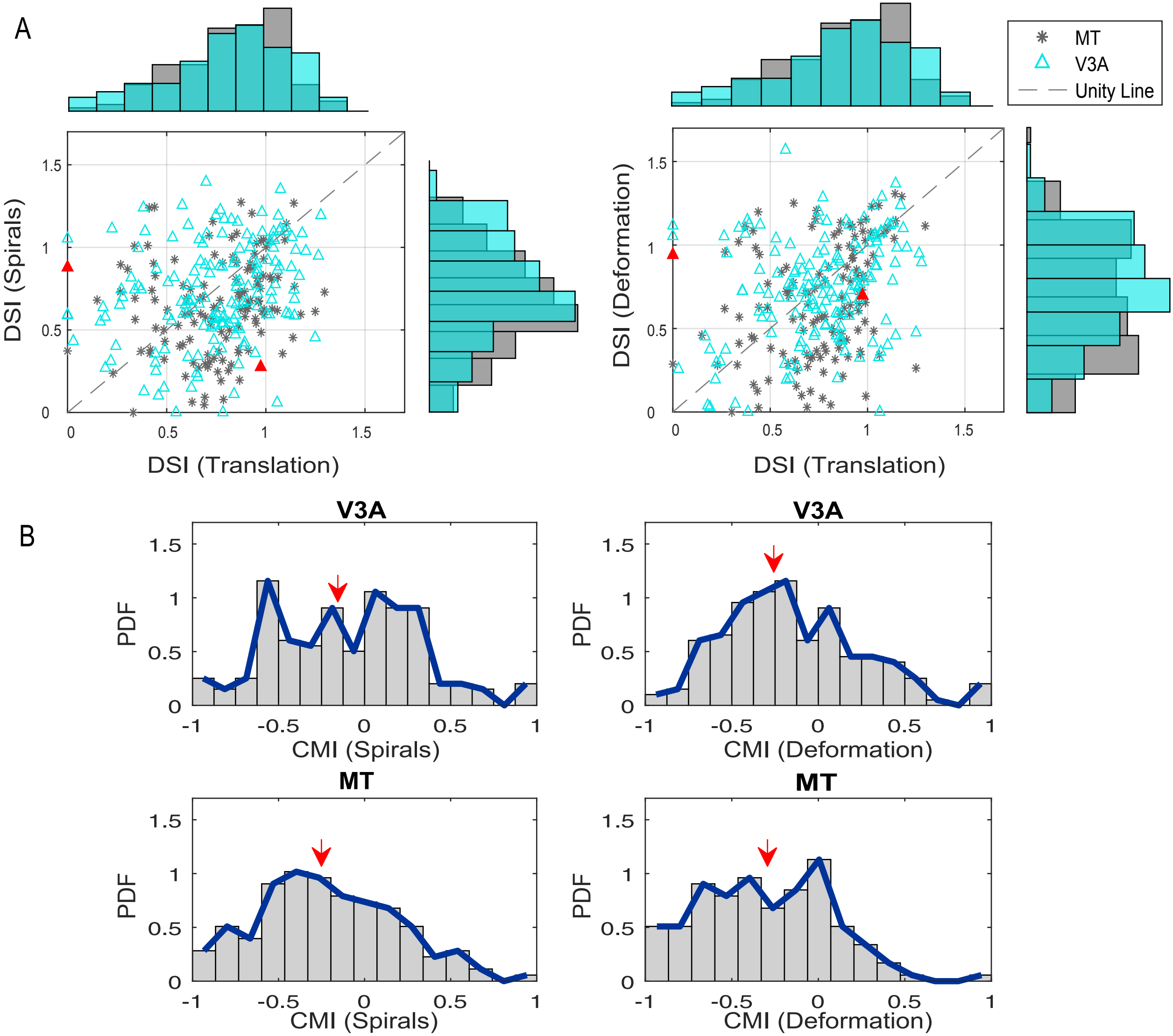

### Figure 7-2

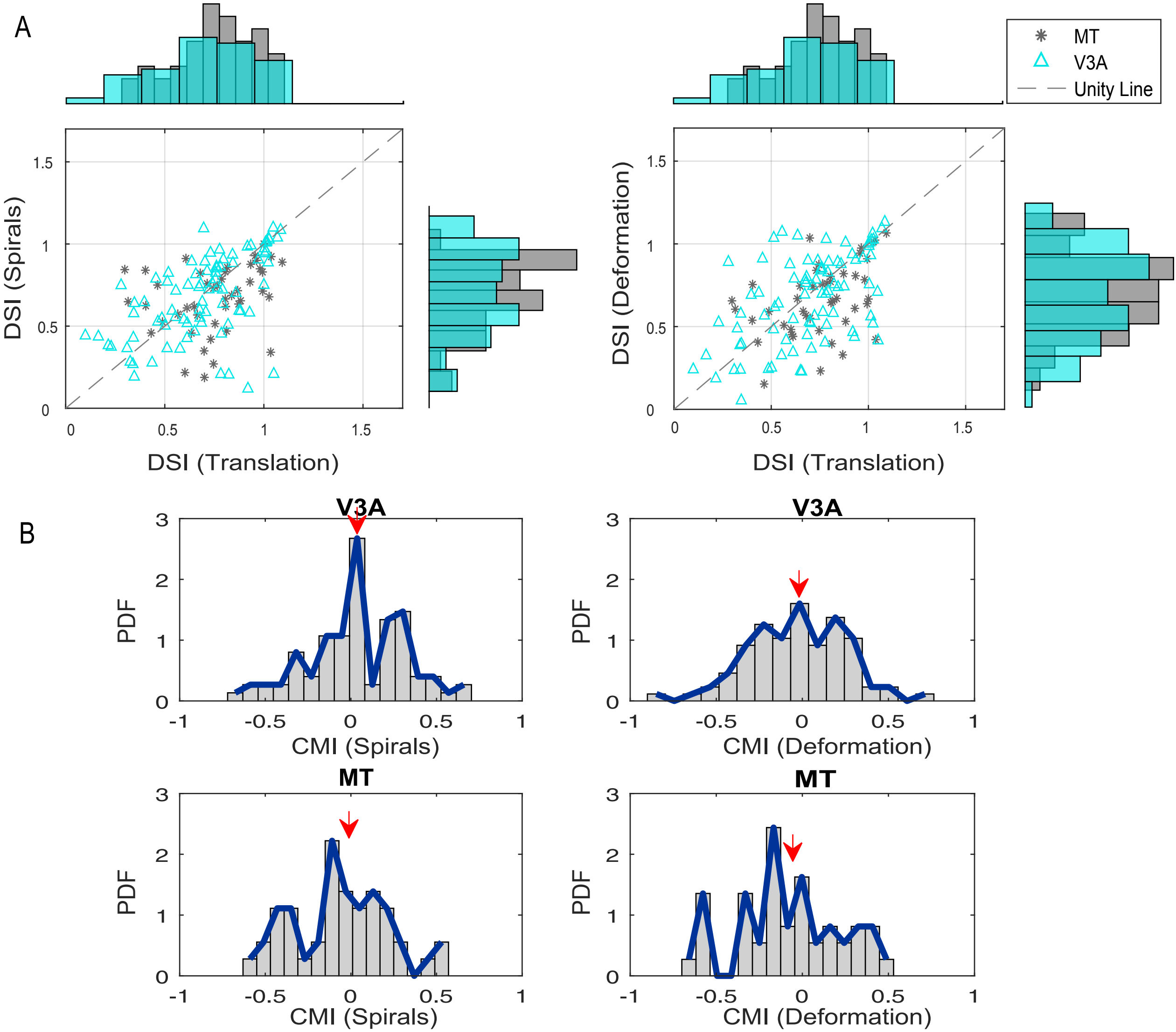
